# Learning to infer transitively: serial ordering on a mental line in premotor cortex

**DOI:** 10.1101/2024.10.29.620924

**Authors:** Sofia Raglio, Gabriele Di Antonio, Emiliano Brunamonti, Stefano Ferraina, Maurizio Mattia

## Abstract

Transitive inference (TI) is a form of deductive reasoning that allows to infer unknown relationships among premises. It is hypothesized that this cognitive task is accomplished by mapping stimuli onto a linear workspace, referred to as the ‘mental line,’ based on their arbitrarily assigned ranks. However, open questions remain: does this mental line have a neural correlate, and if so, where and how is it represented and learned in the brain? In this study, we investigate the role of monkeys’ dorsal premotor cortex (PMd) in encoding the hypothesized mental line during the acquisition of item relationships. Our findings provide evidence that the TI task can be solved through a linear transformation of the neural representations of arbitrarily ranked items. We show that PMd multi-unit activity organizes along a theoretically informed direction, implementing a geometrical solution that effectively explains animal behavior. Our results suggest that the premotor cortex plays a crucial role in integrating item representations into a ‘geometric mental line,’ where the symbolic distance (i.e., rank difference) between items influences the related motor decisions. Furthermore, we observe an ongoing learning process characterized by a rotation of this mental line, which aligns to the linear manifold where motor plan unfolds. This elucidates a cortical optimization strategy based on the statistical structure of the task.

## Introduction

Serial reasoning is fundamental to many aspects of our daily lives, even in the simplest tasks. Consider, for example, the process of preparing a sandwich: while it may seem straight-forward, it involves an intricate sequence of decisions. We start by slicing the bread, then washing a lettuce leaf, and finally cooking a hamburger. Each step is part of a well-defined sequence that guides us toward our desired outcome. Achieving complex goals requires us to assign a specific serial order to our planned actions. As Karl Lashley hypothesized over 70 years ago^1^, this implies that the neural representation of the sequential items in behavior must be encoded before the action begins. The predicted representation of such serial ordering in cortical activity has received substantial confirmation over time^2–5^.

The ability to arbitrarily assign a serial order to abstract items or actions has then to be deeply rooted in our brain and can be viewed as a fundamental computational primitive to be reused across various cognitive functions. A good example of this is transitive inference (TI)^6,7^, a form of deductive reasoning that enables individuals to derive relationships between elements that have never been explicitly compared. For instance, if item A is related to item B, and item B is related to item C, transitive inference allows to deduce the relationship between items A and C, ultimately revealing the ordered sequence of items: A, B, and C. This type of reasoning, known as ordinal knowledge, forms the basis for higher cognitive functions, serving as a fundamental component for algebraic and linguistic computations, as well as hierarchical thinking^7–9^. The transitive inference task has been successfully solved by numerous animal species, including humans, rodents, fish, and non-human primates^10–14^.

Numerous models have been proposed to explain how such a variety of animals can successfully solve the transitive inference (TI) task. Several studies suggest that the presented stimuli are integrated independently within a spatial mental workspace, where each item rank determines its position on a mental line^15–26^. In particular, the ‘geometric mental line’ (GML) model^24^ posits that this mental line is a linear combination of the internal representations of the items, weighted by a monotonic function of their rank. In this framework, the task can be solved by projecting the internal representations of the presented items onto the direction of this line. The symbolic manipulation of disentangled item representations has emerged as a reliable hypothesis for understanding how the brain solves serial learning tasks^27^. This strategy is computationally more efficient than encoding all possible relationships between items and offers significant advantages in terms of generalization and versatility^28,29^. Despite these interdisciplinary efforts, the question of whether this mental line can be encoded in the brain and how it can be learned remains unresolved.

In the outlined framework, the premotor cortex holds significant potential for executing the neural computations necessary to solve the TI task. This region is crucial for decision-making and processing complex information, like those involved in arbitrary sensory-motor transformations^30–33^. Moreover, a growing body of literature demonstrates that the premotor cortex encodes information about the serial order of planned actions to be executed sequentially^4,5^. Previous research studies shown that this brain area is actively involved in logical problem-solving tasks, such as those required in the TI task. Notably, neural activity at the single-unit level in the premotor cortex is modulated by task difficulty in a way that parallels behavioral performance^34^. This relationship is further supported by findings indicating that the premotor cortex functions as a downstream node within a network of areas engaged during the TI task, establishing a privileged interaction with the prefrontal cortex^35^. Given these insights, the activity of the premotor cortex emerges as an ideal candidate for implementing the GML solution, where the ordinal rank of abstract items is encoded within the linear readout of this dynamical system.

In this study, we investigate the role of the dorsal premotor (PMd) cortex of monkeys solving the transitive inference task. We propose that this area is a strong candidate for implementing the geometric mental line (GML) as a potential mental representation for serial reasoning. As a crucial first step, we demonstrate that the PMd cortex encodes task-relevant information, including item identities and their positions on the screen, in addition to serving as a neural correlate for motor decisions. Next, we utilize decoding vectors for each item to construct a data-derived GML based on PMd activity. Our findings reveal that this representation effectively solves the transitive inference task by ordering items according to their ranks, thereby enabling predictions about animal behavior. Finally, we analyze the learning processes occurring during the different phases of the task, illustrating how the animal optimizes its computational strategy by realigning the geometric mental line towards the linear manifold decoding motor decisions.

## Results

### Transitive inference in animals and model

Two macaque monkeys are instructed to perform a transitive inference (TI) task (Fig. 1a, see Methods). In the initial ‘learning phase,’ pairs of abstract visual stimuli (arbitrarily labeled as ‘A’, ‘B’, etc.) were presented, and animals learned to reach the item with highest rank and associated to reward delivery by trial and error. Each session involved a list of seven arbitrarily ranked items, with pairs at symbolic distance SD = 1 (i.e., adjacent items in the list, Fig. 1b). In the successive ‘test phase’, all possible pairs (SD ≥ 1) were randomly presented to assess the animals’ ability to generalize, i.e., to infer transitively the order of items never seen together before. During the experiment^34^, each monkey participated to different TI sessions using series of 6 or 7 items. Here we are reporting data from both animals in the two sessions with 7 items. Two primary behavioral patterns were typically observed in this task^36–38^. The first is the symbolic distance effect (SDE) where the response accuracy (fraction of correct decisions) increases with SD, i.e., with the absolute difference between item ranks in the presented pair (e.g., SD = 1 for ‘AB’ and SD = 3 for ‘BD’). The second is the serial position effect (SPE) which is characterized by the highest accuracy observed in pairs that included the two terminal items.

**Fig. 1.**
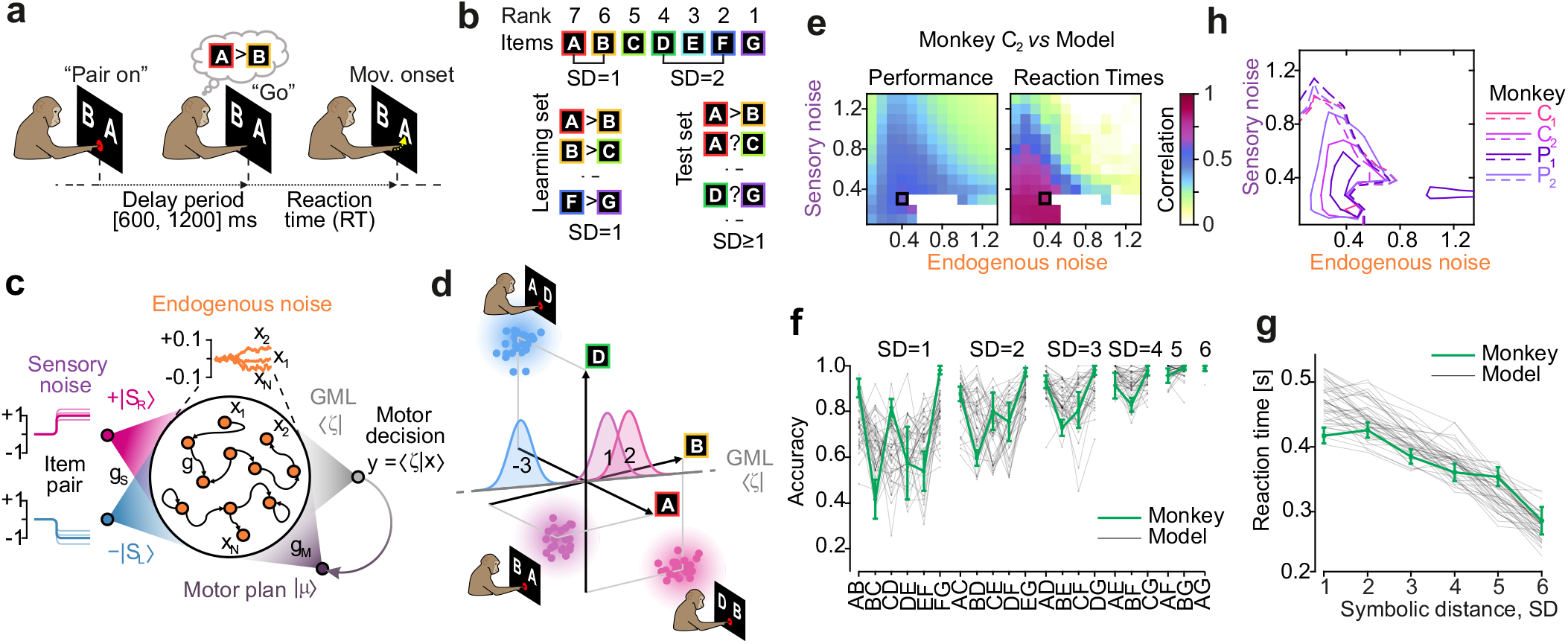
Monkeys and model solve transitive inference with similar performance. **a**, Transitive inference task on a sequence of 7 items. Items are abstract visual stimuli presented in pairs. The monkey has to learn reaching the item on the screen with the highest rank after a random delay ([0.6, 1.2] s) from the “Go” (red cue offset). Reaction times (RTs), times of movement onset from Go. **b**, In the learning phase, only pairs of items with adjacent rank are presented (symbolic distance SD = 1). All pairs (SD≥ 1) are presented in a successive test phase. Each pair is presented in 10 randomly selected trials. Only correct choices are rewarded. **c**, Same task is performed by a recurrent neural network (RNN) with a learned rank-1 feedback with strength *g*_M_ to encode motor plan and output decision with related reaction time^24^. RNN units have stochastic dynamics due to an endogenous noise. The pair of items seen in a trial is modeled as a two-channel input (left and right item) with strength *g*_S_. Items elicit currents drifting RNN activity into independent directions of its state space, randomly perturbed from trial to trial by a sensory noise. **d**, Each item pair is associated to the difference of their representations (axes A, B and D). Inter-trial variability leads to a cloud of RNN states whose projections onto a linear manifold – the geometric mental line (GML) – determine the output decision of the network. The learned GML is given by a suited linear combination of the item representations^24^. Positive (negative) projections determine the decision to reach the right (left) item. **e**, Sensory (*σ*_S_) and endogenous (*σ*_E_) noise modulate behavioral performances and reaction times of RNNs. Spanned intensities of noise can lead to RNN unable to take a decision (white regions). Correlations between experimental accuracies (f, fraction of correct responses) and RTs (g), and those from simulated RNNs determine the model goodness. Black square, RNN best matching with monkey behavior (*σ*_S_ = 0.3, *σ*_E_ = 0.4). **f**, Monkey accuracies (green) in the test phase averaged across sessions (*n* = 2) and animals (*n* = 2) with standard error mean (SEM, error bars). Gray traces, accuracies in RNN with optimal parameters in (e). Only 42 out 100 network simulations are shown; those with response rate as in monkeys (about 70%). **g**, Reaction times grouped by SD for optimal RNNs and monkeys as in (f). **h**, Contour lines at similarity 0.5 in (e) for the 4 analyzed sessions; solid (dashed) contours for performance (RTs).

Various models^12,22^ have explained these effects as either reward-related phenomena or as the consequence of mapping items onto an abstract ‘mental line.’ In this last framework, items with less overlapping representations are associated with higher performance. Single-layer^21,39^ and recurrent neural networks (RNNs)^24–26^ solve the TI task with delta rule learning^40,41^, which is equivalent to classical conditioning. RNNs can also simultaneously encode both rank representations and motor outputs^24^ (see Fig. 1c). However, this simultaneous access to shared computational resources may result in detrimental interference^24^. Specifically, the decision to move to right (left) coded with the output *y* = +1 (–1), is implemented by forcing the network activity to unfold along a specific low-dimensional manifold (the motor plan vector | µ ⟩), as previously reported in PMd^42,43^. We recently demonstrated that under these conditions, RNNs can learn a specific geometrical representation of item ranks, effectively mitigating the potential interference mentioned above. This ‘geometric mental line’ (GML), represented the vector ⟨*ζ* | in Fig. 1d, is given by the linear combination 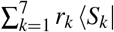 of item representations *S*_*k*_ weighted with the assigned rank *r*_*k*_ (from 7 to 1). The network activity |*x*⟩ – the vector of firing rates *x*_*k*_ of the *N* RNN units – elicited by the input item representations is projected onto the 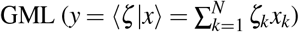. The projection *y* internally drives the network state along the motor plan manifold^24^. When a pair of items is presented, the elicited activity is |*x*⟩ = |*S*_right_ ⟩– |*S*_left_⟩, and its projection onto the GML is the difference of their ranks, which corresponds to the symbolic distance: *y* = SD (Fig. 1d).

In this GML model, both the SDE and the SPE naturally arise from two sources of noise: i) sensory noise, which accounts for the trial-by-trial variability of item representations (illustrated as clouds in Fig. 1d), and ii) endogenous noise, which reflects the intrinsic stochastic dynamics of the firing rates *x*_*k*_ (see Methods). Varying the intensities *σ*_S_ and *σ*_E_ of these two types of noise results in RNNs exhibiting different degrees of similarities to monkeys’ performance (Fig. 1e). Notably, the accuracy of responses for each item pair (Fig. 1f) and the reaction times per symbolic distance (Fig. 1g) during the test phase are best fitted by GML models that share the same set of parameters across sessions and animals (Fig. 1h).

### PMd encodes decision and task difficulty

Given that the GML model accurately mimics animal behavior, a crucial question arises: does the cortical activity in the monkey performing the TI task adopt the same geometrical solution as theoretically predicted? To address this question and to understand what is the role of premotor areas in solving the TI task, we examined the multi-unit activity (MUA) from intra-cortical multi-electrode recordings in the left PMd cortex (Fig. 2a, see Methods). According to previous studies focusing on MUA^44,45^, we found motor plans associated to the decision to reach left or right items well encoded in this area (Fig. 2a). This is apparent from the condition-dependent divergence of the MUA observed in the channels that significantly contribute to the output of the corresponding linear decoder. This finding holds true at both single-channel and single-trial levels (Fig. 2b). Notably, the population code is relatively sparse yet stable across sessions involving the same monkey (Supplementary Fig. 2). In Fig. 1c), we observe that this motor decision matures during the delay period before the ‘Go’ signal^34^ (see also Fig. 1a). The encoded plans are decoded with increasing performances over time, displaying asymptotic (plateau) performances closely correlated with task difficulty, as shown by grouping trials by SD. Task difficulty is greater for smaller SD, resulting in a slower rise (i.e., smaller steepness) of the decoding capacity. Similar trends have been observed in the GML model^24^ and previously shown for single-unit activities in the same experimental setting^34^.

**Fig. 2.**
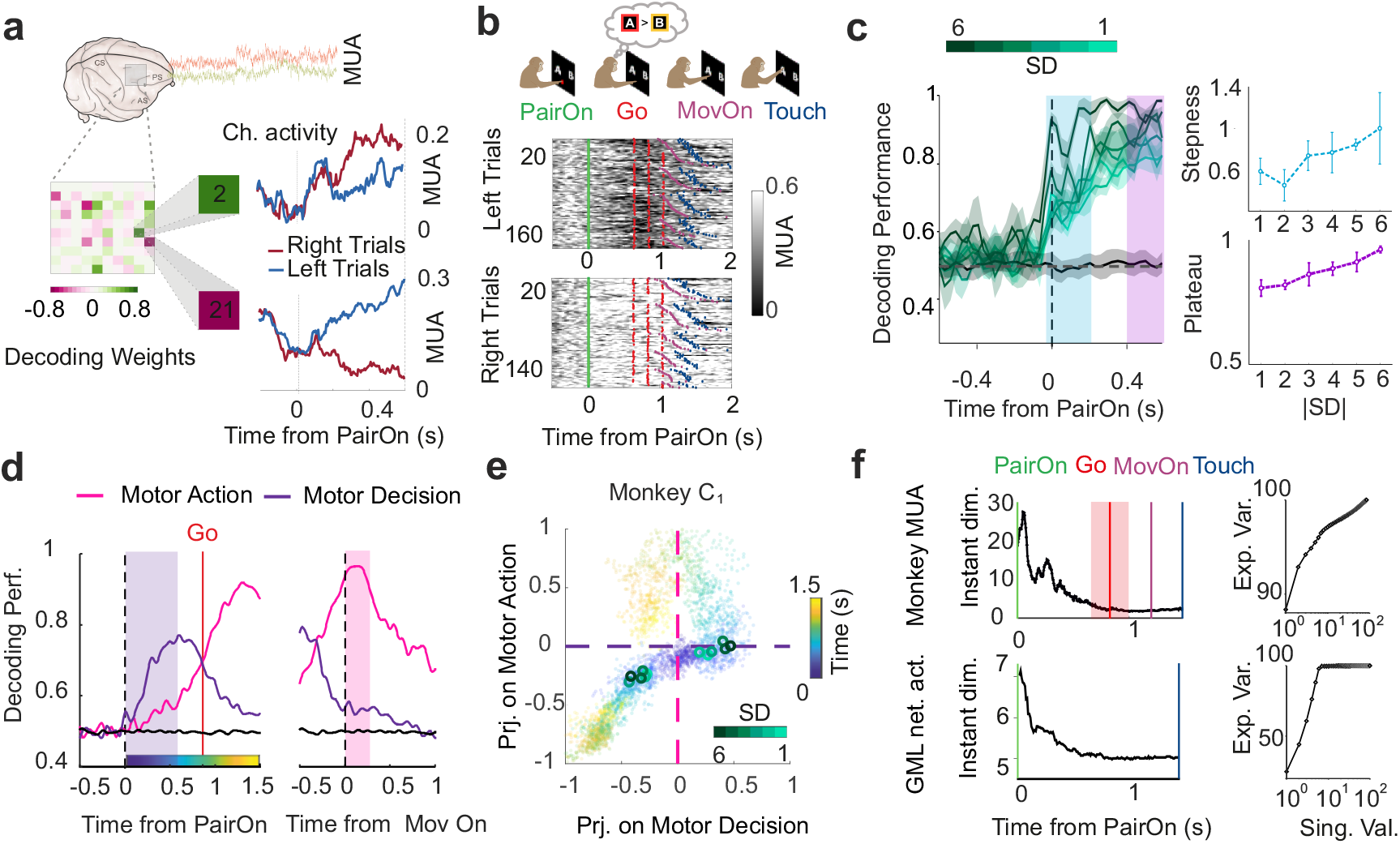
PMd activity predicts animal decision and task difficulty. **a**, Multi-unit activity (MUA) is extracted from unfiltered field potentials recorded by the micro-electrodes arrays (MEA) chronically-implanted in the dorsal premotor cortex (PMd) of two macaque monkeys. Animal decision is linearly decoded from the MUAs of the 96 MEA channels during the delay period. Left matrix, weights of the decoder, i.e., relevance of MEA channels in decoding the motor decision. MUAs from two sample channels averaged across conditions (reaching left or right items) during the test phase. **b**, Raster plots of MUA (test phase, monkey *C*, session 1– *C*_1_) of a sample channel grouping trials for left and right condition. Trials sorted by symbolic distance, and within each SD by reaction times. Markers represent relevant events: green, ‘Pair on’; red, ‘Go’; magenta, ‘Mov. on’; blue, ‘Touch’. **c**, Left, average decoding performances (Area under the ROC curve) across sessions for motor decision divided by SD. Black baseline, decoding on shuffled data. Right, steepness in the rise of performances (slope of the linear fit in [–0.05, 0.2] s after ‘Pair on’) and plateau activity (mean MUA in the interval [0.4, 0.6] s). Both quantities are significantly increasing for increasing SD (slope, *p* = 0.03; plateau, *p* = 0.0001). **d**, Average decoding accuracy (across *n* = 4 sessions) of motor decision and overt movement from MUA (decision and action decoders are trained on the 0.6s and 0.2s following ‘Pair on’ and ‘Mov. on’, respectively). **e**, Projections of the MUA on the (decision, action) decoding axes in the 1.5 s post-‘Pair on’. Time colored according to the color bar in (d); session C_1_. Circles, projections averaged by SD in the post-‘Pair on’ interval [0.2, 0.6]s. **g**, Instantaneous dimensionality of the MUAs in the MEA (top) and of the GML model activity (bottom) estimated from the singular value decomposition (SVD) (see Methods). Right, explained variance.

Movement execution is encoded in the motor cortices as distinct yet complementary information relative to motor plans. This encoding arises from the confinement of neural activity within a low-dimensional latent space characterized by two main orthogonal axes: one representing the decision value and the other representing the reaching movement^42,43,46^. In this context, these two latent dimensions correspond to the decoding vectors for motor decisions and movement execution. Over time, motor plans develop just after ‘Pair on’ signal, eventually giving way to action-related activity following the ‘Go’ signal (Fig. 2d). Within the associated latent state plane, neural trajectories unfold along opposing branches (Fig. 2e). Initially, they drift parallel to the decision axis before shifting toward the action axis, resulting in average projection values that are separated by SD at the end of the delay period.

It is important to note that there is no SD-dependent modulation of decoding performance for motor actions (Fig. 2), which aligns with the hypothesis that these actions have stereotyped neural correlates, as left and right reaching movements remain consistent across trials. Consequently, the delay period is when the premotor cortex can potentially contribute to the neural processing of the TI task. This interval exhibits the highest variability across trials in terms of environmental stimuli. During the test phase, there are 42 possible conditions: 21 pairs presented with the target item on either the right or left. This variability is effectively represented by the dimensionality of the state space explored by the neural activity being probed (Fig. 2f). The highest dimensionality is reached within the first 200 ms following the ‘Pair on’ signal, after which it decreases as the motor plan matures. A dimensionality compression has been observed in PMd cortex also during other motor decision tasks^47,48^. This outcome was predicted by the GML model^24^ and reflects the initial encoding of item representations, which are eventually transformed into a motor plan by drifting cortical activity along the decision axis. In the model, this drift has strength proportional to the projection of the initial activity onto the GML, corresponding to the SD between the items in the presented pair (Fig. 1d). This framework possibly explains both the dimensional reduction observed in Fig. 2f and the SD-related steepness of the performance rising in Fig. 2c.

### PMd encodes both identity and position of items

A key requirement for a cortical network to implement the theoretical GML outlined above is that cortical activity must encode independent representations of the items to be ordered. According to the efficient coding hypothesis, this independence can emerge naturally when input is projected randomly onto the high-dimensional space of network activity. The premotor cortex is expected to have the capability of encoding sequence of items as it is known to receive visual and somatosensory inputs. Accordingly, there is a compelling evidence that PMd cortex operates sensory-motor transformation by actively encoding environmental information along with their serial order^4,30–33^.

We then examined PMd MUAs in the first 600 ms following ‘Pair on’(minimum delay duration), by training 7 independent linear decoders (see Methods). Each decoder was trained on a subset of correct trials to output 1 if the corresponding item was present on the screen and 0 otherwise (Fig. 3a). By projecting the MUAs on the direction pointed out by the decoder weights (namely, the decoder output), it is possible to predict the presented pair of items on average for all the trials (the results for each session separately are shown in Supplementary Fig. 3). This is obtained by picking up the two highest projection values for each condition. Interestingly, decoder output averaged per item are not the same for all of them (orange plots in Fig. 3a). During the learning phase terminal items are better decoded, resulting in a U-shaped performance reminiscent of the serial position effect (SPE). During the test phase the two items with highest rank are better decoded, showing what is known as the ‘magnitude effect’^7,50^ associated with the most rewarded items in the list.

**Fig. 3.**
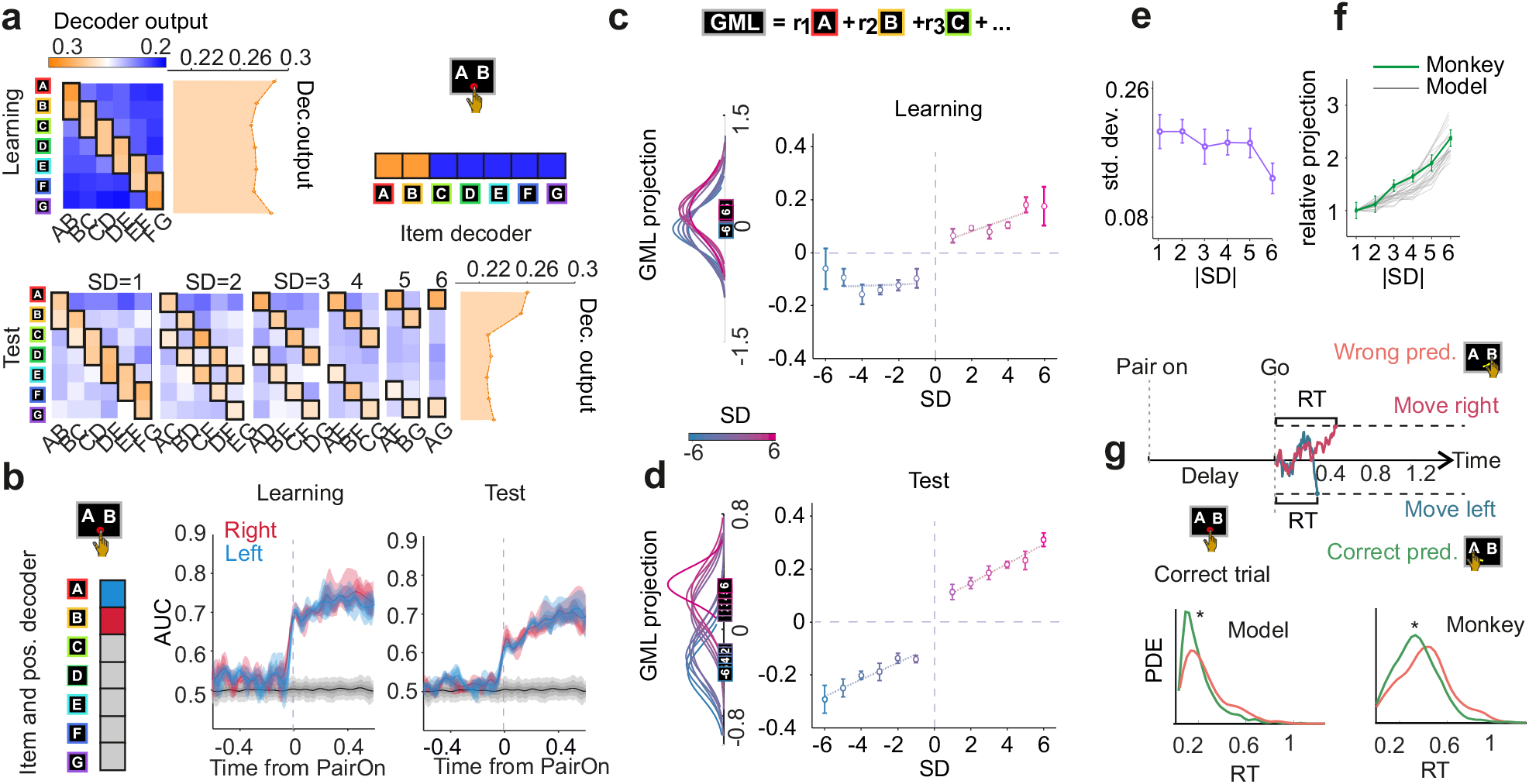
Theoretical GML from PMd represention of items solves the TI task. **a**, Output of linear decoders trained to identify the presence of a single item from MUAs within 0.6 s after ‘Pair on’(minimum delay), regardless of its position on the screen. Top right, example of decoding output in response to pair (A,B). Left, decoding output for all pairs presented in the learning (top) and test (bottom) phase, averaged across all sessions (*n* = 4). Orange histograms, decoder output for each item, averaged over all trials in which is presented. **b**, Performances (AUC, area under ROC) of MUA decoders trained to identify item presence and position on the screen during learning (center) and test (right) phase (averages across sessions, *n* = 4). Left, ideal decoding output in response to pair (A,B): blue, item on the left; red, item on the right; gray, item not present. Shadings represent deciles from 20% to 80%, illustrating the decoding variability across different items. **c**, Projections of MUAs from test trials onto the GML, computed as 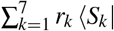, where the *k*-th item has rank *r*_*k*_ and the decoder ⟨*S*_*k*_| (labeled squares in the inset) trained on the learning phase to identify both item and position as in (b). Left, distributions of projections grouped by signed SD, approximated by Gaussians for all trials from *n* = 4 sessions. Mean and SEM across sessions on the right (see Methods for details). **d**, Same as (c) but using item-and-position decoders trained on the test phase as in (b). **e**, Standard deviations of MUA projected onto the GML in (d), grouped by |SD|. **f**, GML projections of MUAs, grouped by |SD |and averaged across *n* = 4 sessions (green). Gray traces, GML projections of the activity |*x*⟩ of the optimal GML models in Fig. 1. Both experimental and model values are normalized to 1 at |SD| = 1. **g**, Top, the GML model decides to move if the projection *y* (colored traces) of its activity onto the GML (Fig. 1c) crosses a threshold (dashed lines for moving right/left), as in classical diffusion models^49^. Bottom, distributions of RTs from the GML model (left) and from monkeys (right) in correct (green) and wrong (red) trials. Experimental trials are from the test phase of all sessions. *, *p* = 2 ×10^−9^ (Wilcoxon rank-sum test). Symbols and error bars in panels (e,f) represent averages and SEMs across sessions (*n* = 4), respectively.

This confirms that PMd cortex encodes differently the items used in the TI task. However, as illustrated in Fig. 1d, the necessary decoding vectors must identify not only the item identity, but also its position on the screen. To achieve this, we trained two linear decoders for each item *S*_*k*_, with vector weights 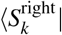 and 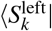 corresponding to the right and left positions, respectively. The representation of item *k* is then given by the vector 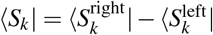. Consequently, the decoding output obtained by projecting the MUAs onto this vector is ideally expected to yield values of +1, –1 or 0 if the item is located on the right, on the left, or is absent, respectively. For example, the output vector for the pair (A,B) is shown in Fig. 3b, where it is also apparent the remarkable performance of these decoders both in the learning and test phase across all sessions and monkeys (Supplementary Fig. 4).

### MUAs projected onto the GML solve the TI task

Having derived the vectors ⟨*S*_*k*_ |from PMd activity representing the items with their screen positions, we can now compute the ‘geometric mental line’. The GML is represented by the vector ⟨*ζ* | of weights of a linear Perceptron. The Perceptron output, given by 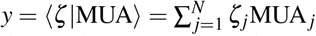, represents the projection of the PMd MUAs extracted from *N* channels onto the GML. Following the same theoretical approach used for the model (Fig. 1c), the GML is defined as the linear combination 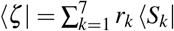. We then projected the MUAs from each trial in the test phase onto the GML computed from the items representations in the learning phase (Fig. 3c). By grouping trials on signed SD (i.e., the rank difference *r*_right_ – *r*_left_), we found that these projection values were predictive of the monkey’s decision to choose either the right (*y* > 0) or left (*y* < 0) item on the screen. Thus, the GML based only on the PMd activity exposed to item pairs with |SD | = 1 is capable to solve the TI task, inferring the right answer also in the test trials with | SD | ≥ 1. Despite this success, average projections were only slightly modulated by the SD of the presented pairs. In contrast, when projecting the same PMd activity onto the GML computed from item representations inferred during the test phase, a SD-dependent modulation emerged (Fig. 3d). As predicted by theory^24^, with increasingly large SD values, projections moved further away from 0, making the motor decision easier and faster. This SD-related modulation was notably steeper when PMd activity was projected onto the GML compared to projections made onto the decoder of motor decision (Supplementary Fig. 5). This suggests that the GML captures a greater amount of task-relevant information. Most importantly, projections of PMd activity onto this GML demonstrated high accuracy in predicting motor decisions made by monkeys at single-trial level, achieving accuracies of 80 ± 2% for correct trials and 60 ± 2% for error trials (mean ± SEM across *n* = 4 sessions, *p* = 0.01 and *p* = 0.01 from chance level, respectively).

Decreasing task difficulty with increasing SD also has a notable effect on the neural data. The distribution of projection values not only shows an increasing average but also a decreasing standard deviation (Fig. 3e). This reduction in variability further decreases the likelihood of errors at larger symbolic distances. The tight link between theory and experiment is also evident when comparing the average projections onto the GML from the experiment data with those measured from the network activity of the GML model (Fig. 3e). In this analysis, averages from Fig. 3d are grouped by |SD| and normalized to 1 at |SD| = 1. Doing the same in the GML model leads to a remarkable overlap, particularly given that the RNNs have been optimized to match the behavioral performance of the monkeys rather than the PMd activity (see Methods). We achieved these results by tuning the endogenous noise that influences the activity of the RNNs, suggesting that the inferred GML model has effectively incorporated the fluctuations characteristic of PMd activity.

In addition to assessing behavioral accuracy, we also evaluated the model’s prediction regarding reaction times (RTs). Within the theoretical framework we propose, the projection *y* of the MUAs onto the inferred GML can be interpreted as the drift in classical diffusion/race models, determining in turn the stochastic accumulation of a decision value^49,51^. In the race model, the decision to move right or left occurs when a corresponding threshold is reached (Fig. 3g-top). Consequently, longer reaction times should be observed in trials where the sign of *y* contradicts the decision made, applicable to both correct and error trials. Specifically, reaching a decision threshold that opposes the direction indicated by *y* is expected to result in prolonged wandering times. Experimental RT distributions from trials where the MUA projection onto the GML fails to predict the decision support this expectation (Fig. 3g-bottom). This aligns with findings from simulations of the GML model. In both scenarios, RTs were significantly slower in trials associated with an incorrectly predicted decision.

In conclusion, our theoretical GML effectively addresses the TI task by identifying the direction in the neuronal state space that accurately sorts internal item representations. Remarkably, this direction can predict trial failures, even when the GML has not been inferred from error trials. Furthermore, model failures correlate with the animal’s reaction time, strengthening the connection between the GML model implementation in PMd and the animal behavior. By optimizing our model based on behavioral data, we thus not only predicted the neural correlates but also uncovered unexpected features in animal behavior, creating a model-experiment virtuous cycle.

### TI task learning implies a rotation of the GML in PMd

So far, we considered the representations of both items and motor plans as stable over time. However, animal behavior during each experimental session reveals that performance evolves, indicating an ongoing learning process. Focusing on the accuracy of responses for pairs of adjacent items (|SD| = 1), we observe that animals’ performances remain relatively unchanged during both the learning and test phases (Fig. 4a-top, purple). This stability is also apparent in the accuracy of the responses from the motor-decision decoder applied to the PMd activity (Fig. 4a-top, green). Not surprisingly, performances for item pairs with |SD| *>* 1 are higher than those for adjacent items, consistent with the symbolic distance effect (SDE). Notably, an adaptive behavior emerges when examining both animal and decoder performances (Fig. 4a-bottom) as they progress from the learning phase through the first half of test trials (Test_1_) to the second half (Test_2_). The improvement in accuracy for the first two groups is partly due to including all symbolic distances in the test. However, the significant increase in behavioral accuracy from Test_1_ to Test_2_ suggests that the monkeys have developed a better representation of item ordering. In contrast, the motor-decision decoder shows no significant changes in performance, indicating that the corresponding motor plans remain unaffected by this learning process.

**Fig. 4.**
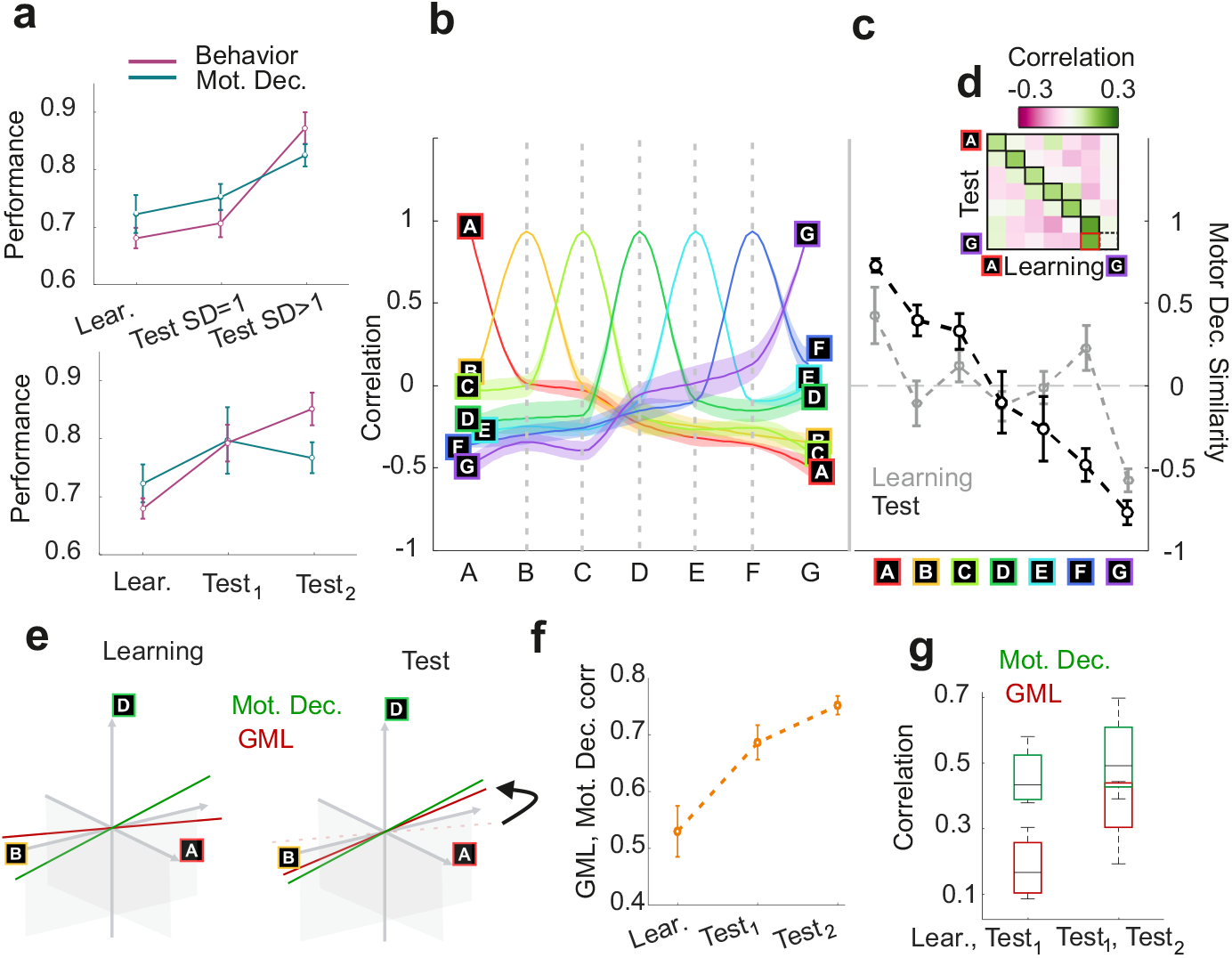
Inferred GML in PMd rotates as a result of learning. **a**, Top, behavioral (purple) and motor-decision decoder (green, as in Fig. 2) performances during learning and test phases. Accuracy of decisions in the test phase is grouped separately for trials with SD = 1 and SD *>* 1. In all panels, mean and SEM (error bars) are over sessions (*n* = 4). Bottom, Behavioral and motor-decision decoder performances during the learning phase, and the first (Test_1_) and second (Test_2_) half of test trials (all SD included). **b**, Correlation between decoding vectors of item identity and position (as in Fig. 3b). Shadings, SEM across sessions (*n* = 4). Curved interpolations are for illustrative purpose only. **c**, Correlations between decoding vectors of items (as in Fig. 3b) and the decoding vector of motor decision, averaged across sessions (*n* = 4) for learning (gray) and test (black). **d**, Correlations between item representations (position-invariant decoding vector as in Fig. 3a) during the learning and test phase. High correlation expected between same items (black frames). Highest wrongly observed (correctly predicted) correlation highlighted in red (black dashed frame). **e**, A sketch illustrating how learning to solve TI task leads to a rotation of the GML inferred from PMd activity (red). Simultaneously, learning does not affect the motor decision axis (green). **f**, Correlation between the decoding vector of motor decision, and the GMLs inferred from the PMd representations of items during the learning phase, and the first and second half of the test phase. Correlations are averaged across sessions (*n* = 4). **g**, Correlation between GMLs inferred from the learning and first test phase, as well as the first and second half of the test phase. Same correlations computed on the decoding vectors of motor decision. Whiskers of box plots represent extreme correlation values; edges are the first and third quartiles; central marks represent the medians.

If motor plans remain stable over time, do other relevant representations in PMd activity adapt in response to changes in behavioral performance? To address this question, we examine the relationships between item representations and their temporal variations. During the test phase, the similarity between decoding vectors of item identity and position (as in Fig. 3b) exhibits a clear pattern of progressive anticorrelations based on their reciprocal distance in the sequence (Fig. 4b). Specifically, item representations during the test phase have a monotonically decreasing correlation (from positive to negative) with the motor-decision decoder, depending on their rank (Fig. 4c). In contrast, such a trend was absent during the learning phase, except for the first and last items in the list (Fig. 4c). Such evidence supports the hypothesis that item representations are influenced by the fraction of times their selection is rewarded.

Collectively, these findings indicate that while learning affects item representations, it does not impact motor plans. Notably, position-invariant representations maintain their similarity across both learning and test phases, with the exception of the last item in the list (Fig. 4d). This picture is perfectly in line with theoretical predictions from the GML model^24^. As learning refines item representations to align with the motor-decision decoder in a rank-dependent manner, the derived GML rotates toward this decoding vector (Fig. 4e), given that it is a linear combination of item representations. We quantified this rotation by measuring the correlation between the inferred GML and the motor-decision decoder, observing an increasing similarity from the learning phase to the second half of the test trials (Fig. 4f). To further confirm that only the inferred GML was rotating, we assessed correlations between the motor-decision decoder during the learning phase, Test_1_, and Test_2_, finding no significant changes (Fig. 4g). In contrast, the GML showed reduced similarity when inferred from the learning phase compared to Test_1_.

In conclusion, as monkeys learn to solve a transitive inference task, the PMd cortex rapidly reorganizes its activity, rotating the geometric representation of the mental line where abstract items are properly ordered. This rotation aligns neural trajectories increasingly closer to the manifold of the latent state space where motor plans unfold, based on the winning statistics of the items.

## Discussion

Transitive inference has long been utilized as a standard task to investigate the implicit learning of sequences and has been considered as a minimal paradigm for testing logical reasoning, generalization, and abstraction. The ability to solve transitive inference task has been widely demonstrated in experimental studies, with successful results observed across a broad range of animal species, including invertebrates^14,52^. However, recent modeling work has been questioning the actual complexity involved in this task solution. In fact, it has been shown that even the simplest linear systems can solve TI task^24,26^. In particular, it has been theoretically predicted that a linear representational geometry, namely the geometric mental line^24^, enables task solution.

Although some work has been done in analyzing neural correlates of transitive inference task solution in non-human primates^34,35,53,54^, these studies have rarely focused on the structure of stimuli representations and their formation during the learning process. In recent years, significant advancements have been made in understanding how representational geometry supports, enables, and shapes behavior during task solutions in non-human primates^27,55–57^ and humans^23,57–61^. These studies typically reveal a strong correspondence between task geometry and the geometry of neural representations across different brain areas.

In the presented work we state that transitive inference task solution is represented in monkeys’ PMd as a projection onto a one-dimensional manifold, i.e., the GML^24^. This theoretically informed direction enables out-of-distribution predictions on error trials and can be interpreted as the input current of a drift-diffusion model of decision making. Remarkably, MUA projections on the GML have the same slope predicted by the RNN model reproducing behavior. In addition to noise levels, a crucial contribution to the slope compression in the model comes from the motor feedback *g*_*M*_: in fact, the analytical derivation of SD values computed from animal behavior in absence of feedback predicts a greater slope (Supplementary Fig. 5b), indicating that part of the information content falls outside the GML subspace, entering the motor decision one. This also happens in the data, as, due to the ongoing learning process, the GML progressively aligns to the motor decision direction, which has lower *SD*– modulation of the projections (Supplementary Fig. 5b) and is stably encoded in premotor cortex over the course of different trials and sessions. This representational drift towards a more efficient strategy is likely driven by changes in the representations and rank values of items, which are updated during the learning process, similarly to what happens in a delta-rule learning of the GML^24^. This is also apparent in the changes in the structure of stimuli correlations, which begin to reflect the statistics of item winning rates. This suggests a statistical learning process during the test phase, which enhances the animals’ performance due to the ‘full-feedback’ paradigm. Similar to other transitive inference task models^26^, this can be viewed as one of the possible shortcuts that make TI task solvable under minimal assumptions^24,25^.

It is important to note that the observed changes in correlations structure are only evident when considering item information conditioned on its position. This suggests that item representations consist of distinct ‘features’: their identity and their rank^62^. The rank, in this framework, can be defined as the correlation between the item vector and the motor output decision, which is highly predictive of behavioral performance. Interestingly, the data-derived GML constructed with the position-dependent item representations (supporting symbolic subtraction) aligns with the motor decision axis, differently from the GML constructed with the position-invariant item representation (supporting symbolic summation, Supplementary Fig. 5e), which remains in a null-potent space with respect to the behavioral decision subspace (Supplementary Fig. 5f). This distinction is likely constrained by the task design. Imagining a different task design, where behavior is sensitive to quantity summation, could potentially lead to the optimization, similarly guided by the learning process, that realigns the position-invariant GML to the decision axis. This would confirm the idea that decoding vectors are reliable item representations^63^ supporting simple algebraic operations^64,65^. All the presented results are investigated in dorsal premotor cortex. This area has already been demonstrated to be relevant in the downstream network involved in TI task solution, following the prefrontal cortex in target encoding^35^ and having a neural activity modulated by different task difficulties, i.e. symbolic distances^34^. While the PFC is typically considered as the primary region for symbolic manipulation of stimuli representations^27,53,55,66^, extensive literature suggests that the PMd plays a crucial role in encoding sensory stimuli, efficiently processing the representations for visuomotor transformation^30,67^, having a more frontal role than traditionally assigned^68^. At the same time, evidence suggests that the prefrontal cortex plays a pivotal role in the planning and execution of motor sequences^3^, making the distinction between the two areas very blurred^69^.

The presented results highlight two interesting features of the PMd activity during TI task solution: i) we found evidence of both position-invariant and position-sensitive encoding, with the latter supporting a natural subtraction operation functional to the decision process, as observed in previous works^33,70^; ii) we found that cortical activity in PMd is modulated by task difficulty, but only during the early phase of stimuli presentation, revealing distinct encoding phases for motor preparation and execution^43,46,71^.

Another intriguing aspect of investigating PMd is observing the ‘readout’ activity of complex brain computations occurring in earlier stages, which aligns well with the linear modeling described in^24^. The fact that we are able to find a linear solution for the task does not imply that the projections of items and pairs on the GML direction should be linear functions of the rank: similarly to what stated in^72^, items less supportive of decision-making (e.g., those in the central part of the sequence, as reflected by behavior) are more orthogonal to the motor decision, and their respective distance is compressed with respect to terminal items, reflecting behavioral performance (Supplementary Fig. 5d).

## Materials and methods

### Subject details

Two male rhesus monkeys (Macaca mulatta), weighing 10 kg (Monkey C) and 9 kg (Monkey P), underwent the experimental procedure with animal care and housing conducted in accordance with European (Directive 2010/63/EU) and Italian (D.L. 26/2014) laws on the use of non-human primates in scientific research.

### Recording apparatus

The neuronal activity of the left dorsal premotor cortex of the two animals was monitored using a 96-channel microelectrode array (Blackrock Microsystems, Salt Lake City, Utah) implanted surgically while they performed a Transitive Inference task. All surgical procedures were performed under general anesthesia (1%–3% isoflurane/O2, to effect) following guidelines for maintaining an aseptic surgical environment. The freeware software package Cortex (https://nimh.nih.gov/) was used to program the presentation of the visual stimuli and the recording of the behavioral responses by a touchscreen (MicroTouch, sampling rate of 200 Hz). The time evolution of the behavioral and neuronal signals were synchronized and recorded by RZ2 TDT system (Tucker-Davis Technologies, Alachua, FL, USA).

### Test stimuli and task design

The two monkeys were trained in identifying the ordinal structure of a 7-item list of abstract black and white fractal images (bitmaps, 16°×16° visual angle) arbitrarily ranked. For each experimental session, a group of 7 images was selected from a dataset of 80 images and then ranked to prevent learning by familiarity. Each experimental trial started with the onset of a starting cue (a red circle with a 13.5°× 13.5° visual angle) in the center of the screen prompting the monkeys to touch it, causing a pair of items to appear on the screen (‘Pair on’), remaining on the screen for a variable time between 600 and 1200 ms. At the end of this waiting period (‘Delay’) the central cue disappeared, signaling the monkey to release the touch (‘Movement onset’) and select the higher-ranking item of the two to receive a liquid reward. The higher-ranking item was randomly presented to the left or the right side of the screen. Each session was divided into three phases (only two of which are considered in the Results): i) a sequential learning phase where the monkey was exposed to pairs of adjacent items in sequence order (e.g., AB, then BC, then CD, etc.) until achieving at least an 80% performance level (Familiarization) in each pair comparison; ii) a learning phase where all adjacent pairs were presented in a randomized order; B (blocks of the Learning phase were presented until the monkeys achieved a proportion of correct trials higher than 60%); iii) a Test phase where the monkey needed to generalize the acquired reciprocal relation between the adjacent items sequence to infer the relationship between all possible pairs of items. In both, Learning and Test phase, each pair was presented the same number of times (20 trials per pair, with 10 trials having the winning item on the right and 10 on the left side of the screen).

### Behavioral data analysis

Behavioral data reported in Figure 1**f**,**g** show the accuracy and reaction times (RTs) for different symbolic distances (SDs). The accuracy is computed as a percentage of trials of the Test session in which the animal answers correctly. The trials that are aborted before the animal touches the wrong item are not included in this computation. The RTs are computed from the onset of the Go signal to the Movement onset. All the trials are considered in this computation, both correct and error ones. The plotted values are averaged across the four sessions, with the corresponding standard error of the mean (SEM). The same computation of accuracy and RTs is used to produce Figure 4**d**,**e**.

### Neural data pre-processing

Multi-unit activity (MUA), depicted in Figure 2**a** and used for all the successive analyses, was extracted by computing the time-varying power spectra from the short-time Fourier transform of the unfiltered extracellularly recorded field potential in 5ms sliding windows, normalized by the average power spectrum across the session in a [200, 1500]*Hz* frequency band. Logarithmically scaled MUAs were smoothed by a moving average (40 ms sliding window). This procedure is the same as in^45,73^. For the analyses we considered 85 out of 96 channels per grid, which were common to all sessions for both animals after anomaly detection of the recorded activity. Off-line analyses were implemented in MATLAB (The MathWorks).

### Neural activity decoding

The linear decoding of the 85-channel multi-unit activity was executed using pseudoinverse operations implemented in MATLAB. The decoding algorithm was trained on 70% of the data and tested on the remaining 30%, which consisted of independent samples of multi-unit activity, with one sample recorded every 50 ms. The logMUA was preprocessed utilizing a 40 ms moving average. The decoding was performed 100 times on different data subsets – an overlapping cross-validation – returning the average performances in terms of accuracy, computed using the optimal threshold between the two classes returned by the AUC (area under the receiving operating characteristic curve) computation. After verifying that the performances on test set are significant with respect to the chance level, we define the representation of each decoded variable as the average value of its decoding vector across 10 realization. In the manuscript results we describe the decoding of different variables: i) the motor decision, ii) the motor action, iii) the symbols and iv) the symbols associated to their position. Each of these variables has some peculiarities in the decoder training, which are specified below.

In the case of motor decision, the decoder is trained on the time interval [0, 600]ms after ‘Pair on’, always before the ‘Go’ signal to avoid any contamination of movement preparation and onset. The target output is +1 when the animal moves right and −1 when it moves left. The decoder is trained on both correct and error trials in order to disentangle information about the winning symbol position and the animal motor decision. We consider as errors only the trials in which the animal chooses the wrong symbol, not those aborted before.

As for the motor decision, in the motor action decoder the target output is ± 1 for the right (left) movement of the animal and both correct and error trials are included in the training.

The considered training interval, instead, in this case is [0, 200]ms after the ‘Movement onset’.

For the symbols position-invariant information, we use 7 decoding vectors, one for each symbol, asking as a target output +1 for the trials in which the symbol is presented on the screen, independently on its position, and 0 when the symbol is not presented. Both the training and the test sets are re-balanced, sub-sampling the majority class, which is always 0 (we are considering one symbol among seven in every decoding, so there are many trials in which that specific symbol is not presented). We only consider correct trials for this decoding, as we do not have a specific hypothesis about why the animal is choosing the wrong symbol and we can not exclude the possibility that the animal has a wrong encoding of symbols during the error trials.

For the symbols and position information, for each of the 7 items we want a decoding vector which has as output +1 if that symbol is present on the screen and it is placed in the right side, −1 if it is present and placed on the left side, and 0 if the symbol is not on the screen. We obtain a decoder equivalent to this computing the difference between these two decoders: i) the right-position decoder returning +1 when the symbol is present and placed on the right and 0 in all the other conditions (including when the symbol is present but on the opposite side) and ii) the left-position decoding returning +1 when the symbol is present and placed on the left side of the screen and 0 otherwise. As for the symbols decoding, here we only consider correct trials and re-balance the classes.

In the plots showing the decoding performances in time (Figure 2**c**,**d**, Figure 3**c**) the baseline value is computed performing exactly the decoding described above but with shuffled labels. The already computed vector is tested for each time point in a specified interval, and the AUC value (Figure 2**c**, Figure 3**c**) or the accuracy (Figure 2**d**) is reported. In the specific case of Figure 2**c** a different decoding vector is computed for each SD. In Figure 4**e** above, the reported decoding performances refer to the motor decision decoder, trained only on Learning, Test pairs with *SD* = 1 and Test pairs with *SD* > 1, computing across trials the accuracy of its average projection value over all the considered time interval ([0 600]ms from Pair on). The same holds for the plot below, where the decoder was, instead, trained during Learning, the first half of Test trials and the second half of Test trials.

All the correlations between the variables representations (i.e. decoding vectors) are Pearson correlations (Figure 3**b**, Figure 4**f**,**g**,**h**). The correlations among symbols and between symbols (Figure 3**b**, Figure 4**f**,**g**) are computed excluding the bias value, while those between GML and decision or among decision vectors (Figure 4**h** and Supplementary Fig. 2, Supplementary Fig. 4) include also the bias value.

### The GML model and its application to data

This work applies the Geometric Mental Line (GML) model^24^ on neural data of monkeys performing a transitive inference task. The GML is a linear combination of symbolic representations, weighted by a monotonic function of their rank, denoted as *f* (*r*_*j*_), where *j* ∈ [1, *M*] and *M* is the number of items in the list. Specifically, it is formulated as follows:

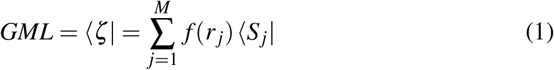

with *S* _*j*_ being the item representation. In the specific context of transitive inference, fixing the same reward value for each pair, it holds that *f* (*r* _*j*_) = *r* _*j*_^24^.

The data-derived GML illustrated in Fig. 3**c-g** and Fig. 4**f**,**g** is computed using the position-sensitive decoding vectors of the stimuli as ⟨*S* _*j*_| in Eq. (1). The coefficients *f* (*r* _*j*_) are determined by presenting all possible pairs of adjacent item decoding vectors (12 pairs) along with the corresponding output decisions (+1 for the target item on the right, −1 for the target on the left) to a pseudoinverse algorithm.

The GML model has been found to be spontaneously implemented by recurrent neural networks performing the same task, thereby providing a digital prototype of brain functioning. In^24^, we proposed a linear RNN characterized by a one-dimensional manifold encoding the motor plan. This exemplifies the premotor cortex of monkeys, known to contribute to motor decision-making in transitive inference task, with neuronal activity modulated by the symbolic distance between item pairs^34^. The recurrent neural network consists of *N* = 100 units with activity states defined as *x* _*j*_(*t*) = |*x*(*t*) _*j*_⟩. The network receives sensory input when a pair of items (|*S*R– *S*_L_⟩, representing right and left symbols) is presented, and an additional motor-related input | µ⟩ is added and modulated by the expected response *y*^′^(*t*), leading to the following linear dynamics:

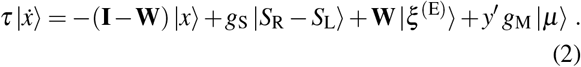

In this equation, *g*_S_ and *g*_M_ modulate the strengths of sensory and motor inputs, respectively. The elements *W*_*jk*_ compose the synaptic matrix **W** ∈ ℝ^*N*×*N*^ sampled randomly from a Gaussian distribution with zero mean and standard deviation 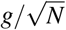.

Each unit receives a current of endogenous noise |*ξ* ^(E)^⟩, uncorrelated in time, whose elements 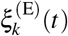 are i.i.d. random variables drawn from a Gaussian distribution with zero mean and an arbitrary variance 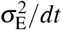. This description aims to model the stochasticity observed in biological neuronal networks operating under conditions of balanced excitation and inhibition^74,75^.

In addition to endogenous noise, sensory noise is incorporated into the model to account for variability in encoding visual stimuli across trials. The effective contribution of a presented pair of items in a generic trial *n* is given by 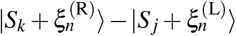, where 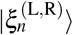 represents a vector of i.i.d. Gaussian values with zero mean and standard deviation *σ*_S_.

Within this framework, the readout weights ⟨*ζ*| defined by Equation (1) enable the network to solve the transitive inference task on average. This RNN serves as the basis for fitting the behavior of two monkeys (Fig. 1**e**,**f**,**g**,**h**) and generating comparisons between digital simulations and real data, as shown in Fig. 2**f** and Fig. 3**f**,**g**.

Each evaluation is based on correlation between animal and network behavior (performance and reaction times) for different noise parameters, considering only networks that respond to more than 70% of trials. The feedback parameter for the RNN is set at *g*_*M*_ = 0.01. When network activity fails to reach a predetermined threshold, it is treated as non-responsive, similarly to trials where an animal does not select a symbol within a fixed time frame during experiments. The number of networks that responded to more than 70% of trials for different noise parameters is reported in Supplementary Fig. 1**a** (the percentage of trials in which animals do not respond is approximately 85%± 5% across sessions). Response rate and occurrence of behavioral effects (SDE for performance and RTs, SPE) in simulation varying noise parameters is reported in Supplementary Fig. 1**b**. In Fig. 1**e** and Supplementary Fig. 1**c**, each value on the grid represents the correlation value between the performance curves (21 performance values, one for each pair as in Fig. 1**f**) of the animal and the network, and between the RTs curves (6 values, one for each symbolic distance as in Fig. 1**f**) of the animal and the network, for varying numbers of networks.

### Dimensionality reduction

The dimensionality time course in Figure 2**g** has been performed as in^76^, computing the SVD of the concatenated trials from the Test phase, centered around the Pair onset and averaged per condition (42 possible pairs). The explained variance per dimension is also reported in the plot. The time course is averaged across conditions and the same computation is applied to the internal activity of the RNN with feedback and fixed noise values (*g*_*M*_ = 0.01, *σ*_*E*_ = 0.4, *σ*_*S*_ = 0.3) performing TI task described above.

## Acknowledgements

Work partially funded by EU H2020 Research and Innovation Program, Grant 945539 (HBP SGA3) and by the Italian National Recovery and Resilience Plan (PNRR),M4C2, NextGenerationEU (Project IR0000011, CUP B51E22000150006, ‘EBRAINS-Italy’) to S.F. and M.M.

---

**Supplementary Fig. 1.**
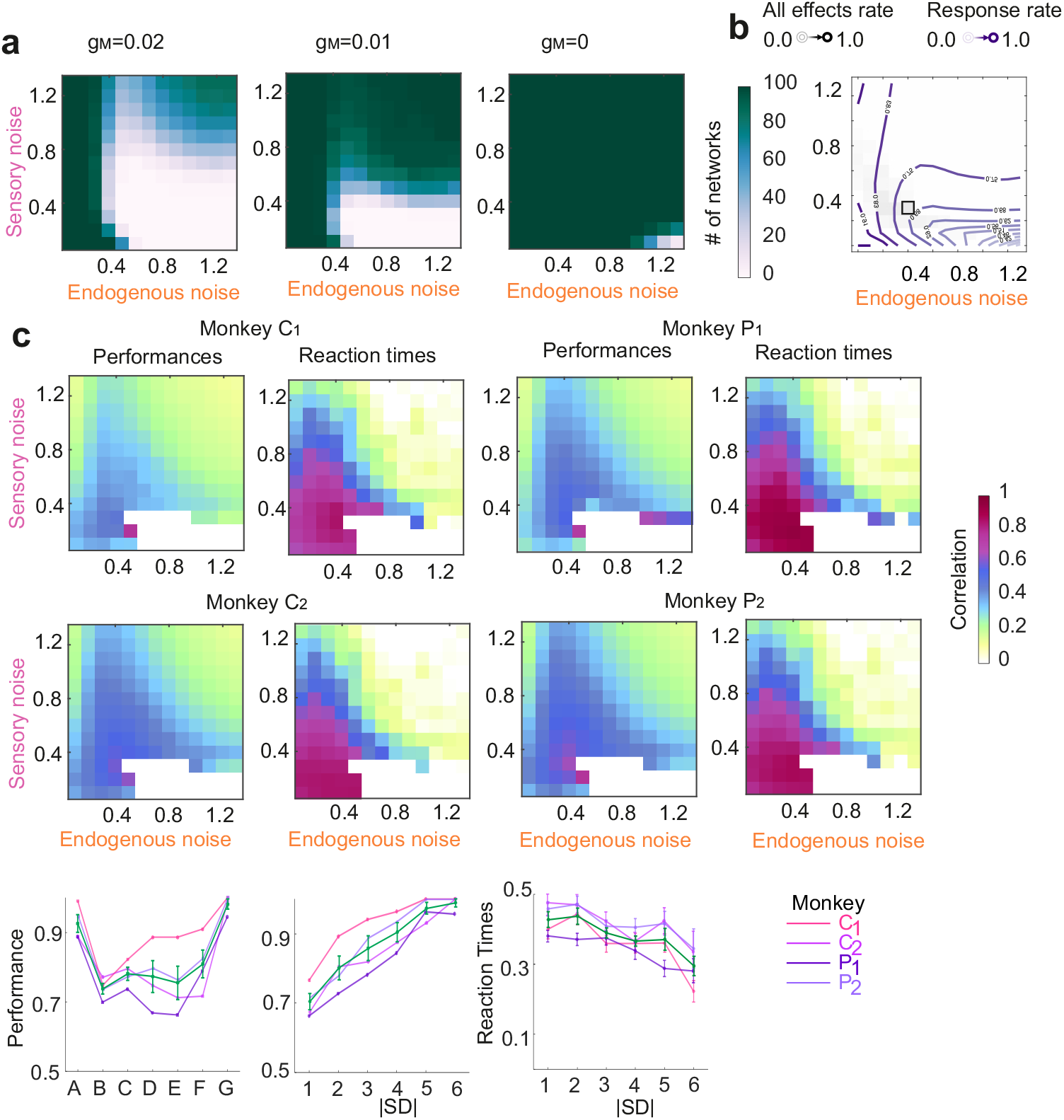
**a**, Varying the feedback intensity (*g*_*M*_ = 0, 0.01, 0.02) the number of networks over 100 tested that reach the response threshold of 0.5 in at least the 70% of the trials depends on the noise intensity (both sensory and endogenous). The stronger the feedback, the lower the number of responsive networks. **b**, Rate of simultaneous occurrence of i) SPE on accuracy and ii) SDE on both accuracy and reaction time on the GML with *g*_*M*_ = 0.01. The contour plot indicates the response rate in the different sensory and endogenous noise parameters region. The black square shows the effects rate and the response rate in the noise parameter chosen in Fig. 1a. **c**, Cosine similarity between each monkey performances and reaction times and the GML model (*g*_*M*_ = 0.01) with different levels of noise. As reported in main Fig. 1d the best fitting area of the different sessions and animals is similar. **d**, From left to right, four sessions performances showing SPE and SDE and reaction times showing SDE with superimposed average value with SEM, as reported in Fig. 1f,g.

**Supplementary Fig. 2.**
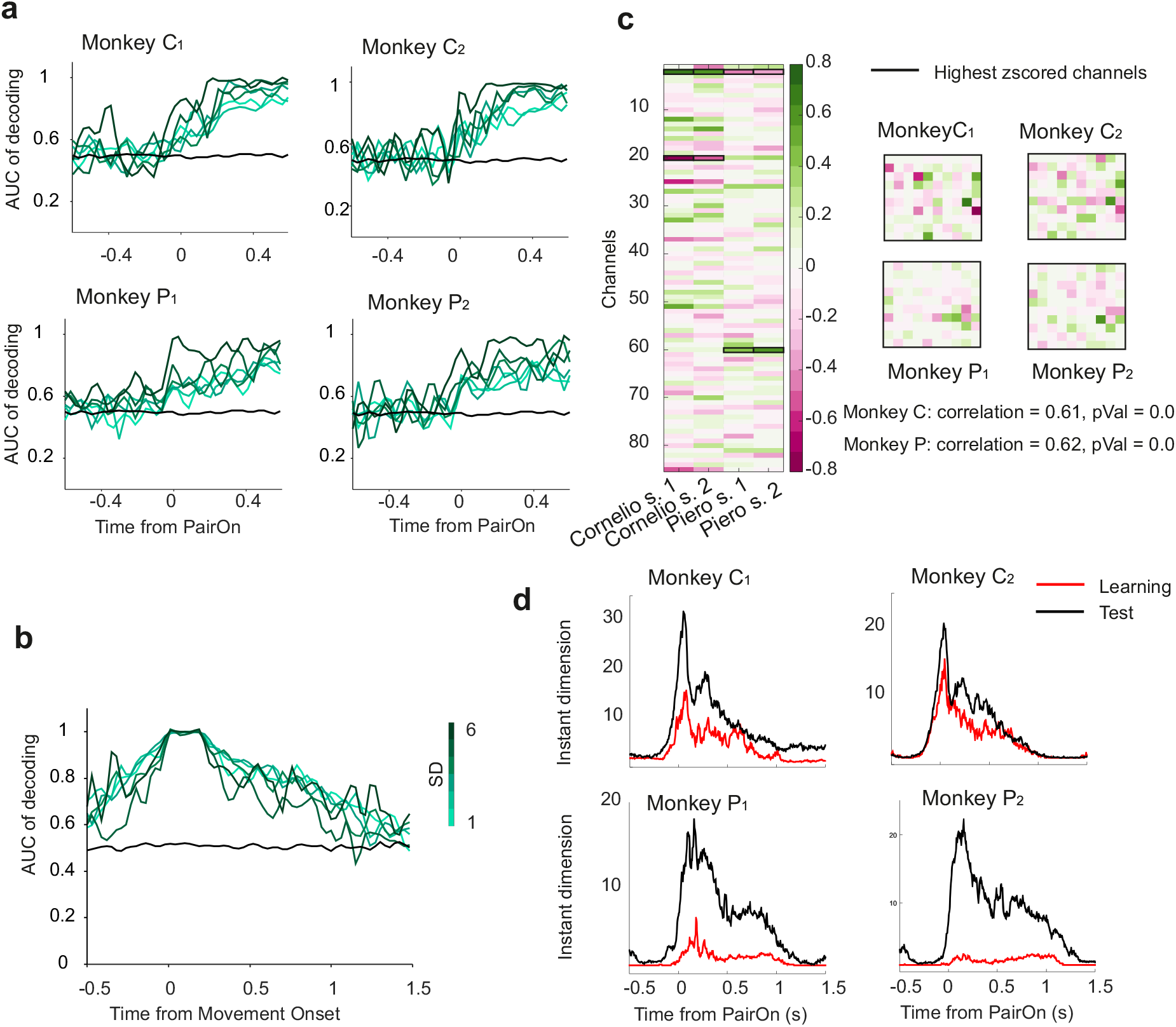
**a**, Decoding performances (AUC) for the motor decision modulated by SD for each of the four sessions. **b**, Decoding performances (AUC) for the motor action are not modulated by SD. The reported plot is averaged across the 4 sessions. **c**, Decoder weights for the four sessions, highlighting the most relevant channels (*zscore* > 2) for each session. Interestingly, the decoder weights patterns are similar in the two sessions of the same animal, meaning that the motor decision representation is stable in different days. Stability of decision encoding for the two different sessions of the same animal reported as correlation and p-value. **d**, Instant dimension for each session comparing the learning and the test phase. The profile is similar but interestingly the dimensionality of learning is smaller than test one. This could be due to a lower complexity given by the presentation of only adjacent pairs (12 conditions compared to the 42 conditions during test).

**Supplementary Fig. 3.**
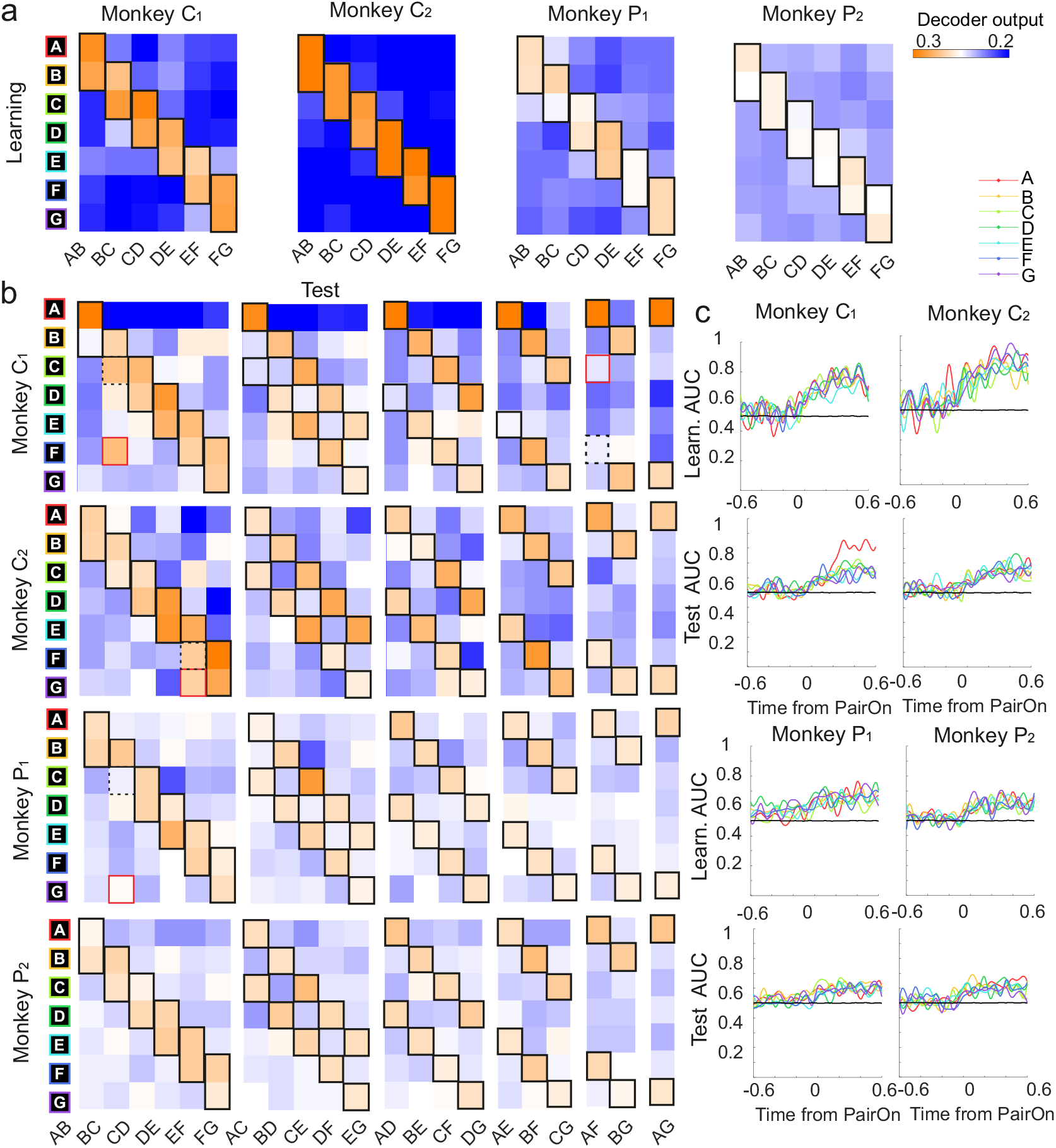
**a**, Projections of the MUA onto the 7 position-invariant decoding vectors for each of the four sessions during learning (average projection across sessions is shown in Fig. 3a). Presented pairs are correctly predicted. **b**, Same as in (a) but for the test phase. Correct predicted items are framed in black. Wrong predicted items are highlighted in red and the correct expected prediction is dotted in black. **c**, Decoding performance (AUC) of the position-invariant decoders in (a), (b) for time steps centered around PairOn. From left to right, the four Learning sessions and the four Test sessions. Different color codes are used for the different symbols decoding performances.

**Supplementary Fig. 4.**
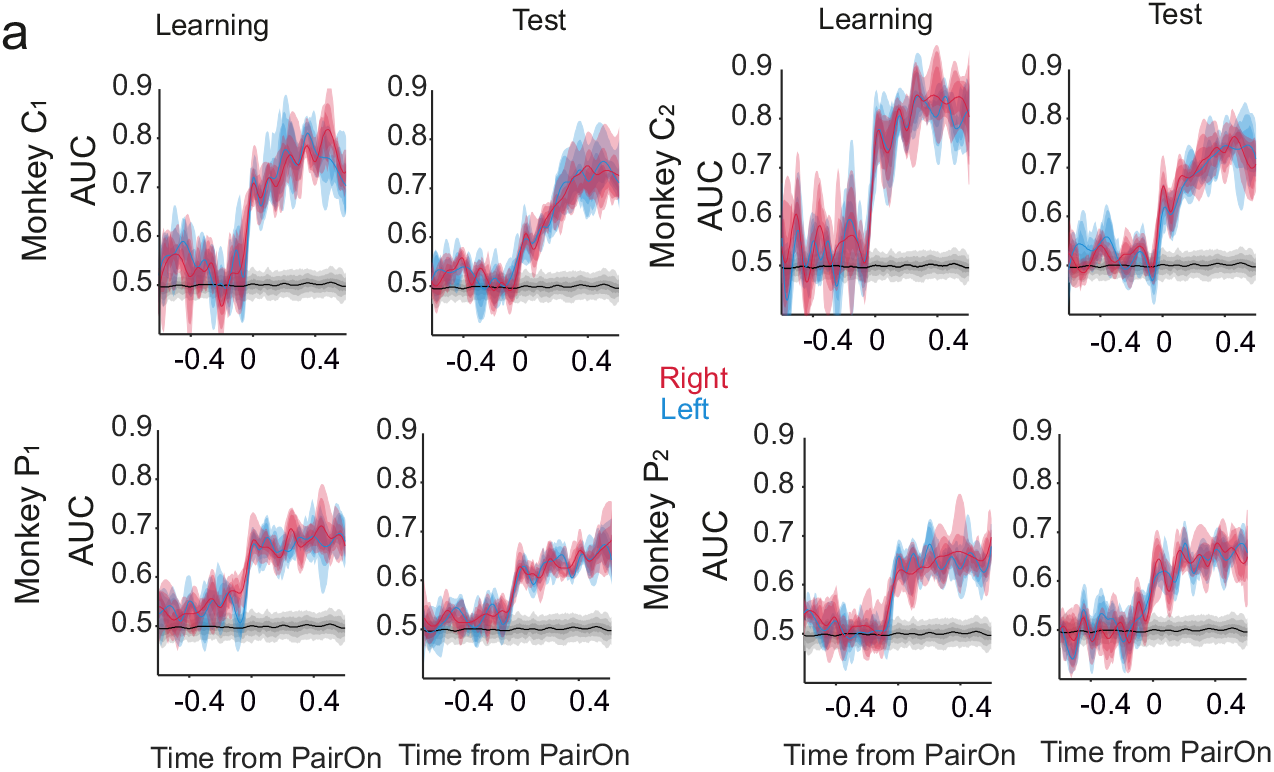
**a**, Learning and test position-sensitive symbols decoder performances (AUC) on the four sessions separately. Average across sessions is shown in Fig. 3b. The shading indicates percentiles from 20 to 80 referring to the variability of decoding across symbols.

**Supplementary Fig. 5.**
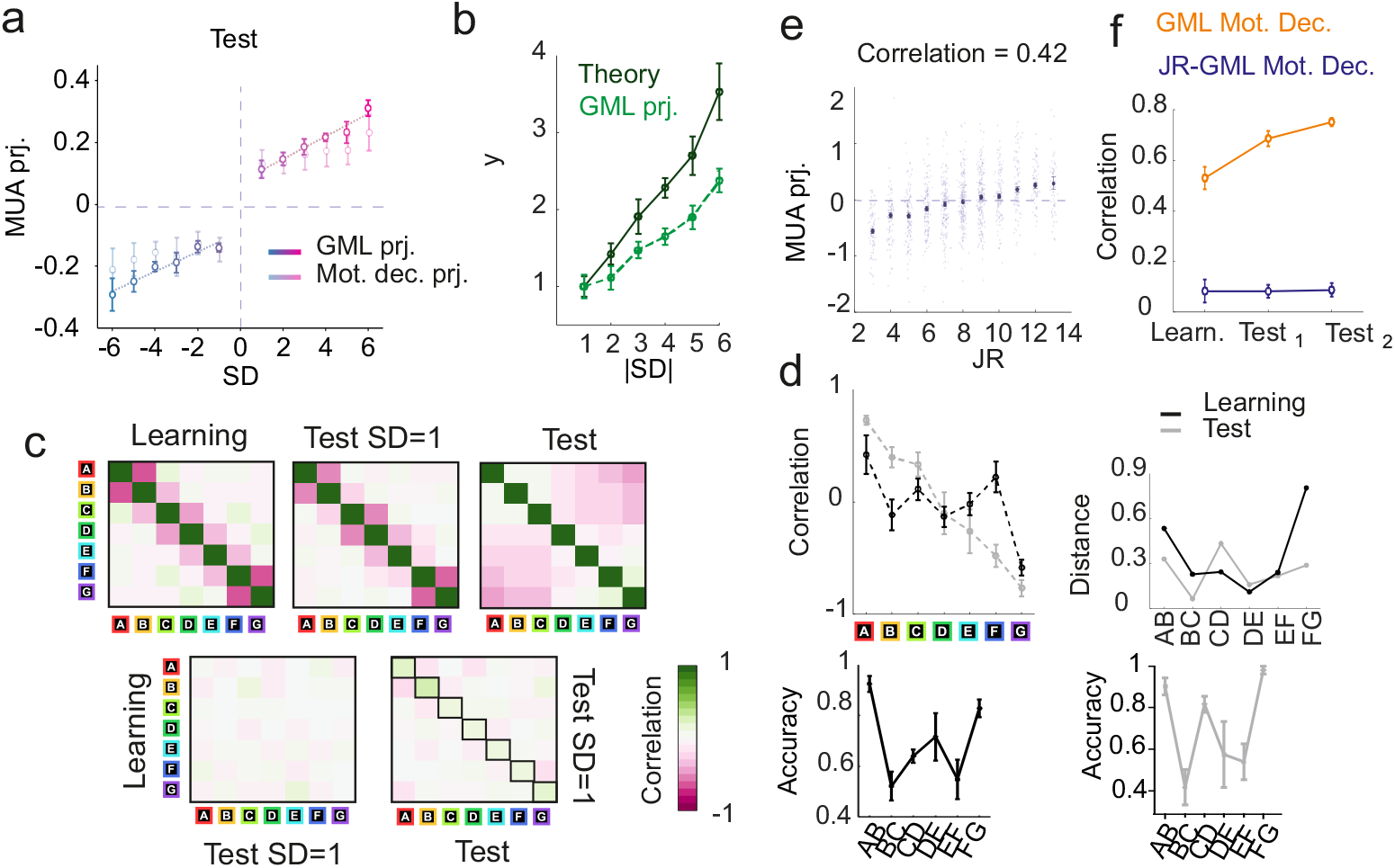
**a**, Comparison between the MUA projections for trials with the same signed SD on the GML and on the motor decision decoder during the test phase, on average across sessions. Both directions are predictive of the animal decision but the GML has a steeper modulation by SD. **b**, Comparison between the GML projections predicted by the theory and the actual projection values as reported in Fig. 3**f**. The theoretical prediction (not taking into account feedback) is computed using the formula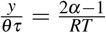, where*y* is the projection value, *θ, τ* are normalization factors indicating the decision threshold and the model time constant, *α* is the accuracy and *RT* the reaction time for every symbolic distance. **c**, Top. Correlation matrix between symbols representations during Learning, Test considering only trials with SD=1, Test. Bottom. Correlation matrix between symbols representations transiting from Learning to Test (*SD* = 1) and from Test (*SD* = 1) to Test. The correlation structure is maintained between Learning and Test (*SD* = 1), while the representations change dramatically. Conversely, the symbols representations are maintained between Test (*SD* = 1) and Test, while the correlation structure changes completely. **d**, Top, Left. Correlation between motor decision and symbols vectors for the learning and the test phase (as in Fig. 4**c**). These values can be considered as a proxy of the symbols ranks. Top, Right. Distance between the correlation values (Top, Left) of adjacent symbols for both learning (Left) and test (Right). The non-constant distance values indicate that symbols are projected with a non-linear function of their ranks. Bottom. Behavioral performances for *SD* = 1 during learning and test on average across 4 sessions. The accuracy is predicted by the distance between the projections (Top, Right). **E** Projections of the MUA for trials divided by joint rank (JR) on the GML built with the position-invariant decoders. There is a significant modulation of the activity tuned by JR parameter (*p* = 0.0). **f**, Correlation between the GML and the motor decision vector during Learning, first half of the test trials, second half of the test trials (as in Fig. 4**f**). Same for the correlation between the GML built with the position-invariant decoders and the motor decision vector.

